# Robust PHP in Adult Hippocampus: Essential Assay Optimizations

**DOI:** 10.64898/2026.03.12.711375

**Authors:** Peter H. Chipman, Richard D. Fetter, Forrest J. Ragozzino, Unghwi Lee, Graeme W. Davis

**Affiliations:** Department of Biochemistry and Biophysics Kavli Institute for Fundamental Neuroscience University of California, San Francisco San Francisco, CA 94158

## Abstract

Presynaptic homeostatic plasticity (PHP) is a potent form of homeostatic plasticity that has been documented at synapses as diverse as the glutamatergic *Drosophila* neuromuscular junction (NMJ), cholinergic mammalian NMJ (including human), and glutamatergic synapses in the mammalian brain. Published experimental evidence in favor of PHP in adult hippocampus and cerebellum includes patch-clamp electrophysiology, presynaptic capacitance measurement, calcium imaging, optical reporters of vesicle release and correlated three-dimensional electron microscopy. These studies are grounded in newly optimized experimental protocols that differ substantively from those typically used to study activity-dependent plasticity in neonatal and juvenile slice preparations. Here, we elaborate and extend our assays and methodologies for the study of PHP in the adult mammalian brain. Our assays are designed to optimize synapse, cell and tissue health and minimize the incorporation of unintended adverse experimental conditions that may interfere with the induction and/or expression of PHP. In addition, we provide benchmark criteria for assessment of cell health, necessary for analysis of PHP and, in so doing, advance our understanding of postsynaptic conditions necessary for PHP induction in the adult brain. Our data underscore why PHP may have been previously overlooked, inclusive of a recent manuscript challenging the robust expression of PHP in the mammalian brain (Dou et al., 2026 BioRxiv [preprint]).

## INTRODUCTION

Homeostatic plasticity (HP) encompasses a suite of compensatory physiological processes that have been shown to counteract perturbations and stabilize neural function ^1–12^. The term presynaptic homeostatic plasticity (PHP) describes an activity-independent phenomenon observed at the neuromuscular junctions (NMJ) of *Drosophila*, mouse and human ^9,13–17^. Recently, three papers pioneered the characterization of PHP at mammalian central synapses in acute brain slice ^18–20^. In brief, at excitatory synapses in both cerebellum and hippocampus, partial antagonism of postsynaptic AMPA receptors using highly selective antagonists (GYKI or perampanel) initiates a rapid (∼30 min) compensatory response that potentiates presynaptic neurotransmitter release and preserves the gain of synaptic transmission at levels observed prior to application of the AMPA receptor antagonist. Additional data demonstrate that PHP, in both hippocampus and cerebellum, can be induced by partial genetic depletion of postsynaptic AMPA receptor abundance, paralleling observations at neuromuscular synapses in *Drosophila*, mouse and human.

In support of current experimental progress studying PHP in the mammalian brain, it should be emphasized that every assay in experimental biology confronts variables that need to be addressed. Some variables are known and controllable, while others inevitably remain unknown. Thus, no single assay, no single measurement, no single technique can be assumed to provide unequivocal evidence of a biological process or mechanism, particularly one as complex as homeostasis. Confidence accrues when multiple, independent assays, techniques and measurements converge on a common conclusion. This is precisely what has been achieved in favor of PHP ^18–20^. Published assays that support the presence of robust PHP expression in adult brain are extensive and diverse including: **1**) Multiple electrophysiological measures were made (EPSCs, IPSCs, estimates of short-term plasticity and vesicle pool dynamics) across two different brain regions (hippocampus ^19,20^ and cerebellum ^18^). **2**) Mechanistic pharmacology was performed including dual-patch clamp recordings providing evidence that postsynaptic NMDARs are necessary for PHP ^20^. **3**) Optical assessment of neurotransmitter release was performed using GluSnFr-based estimates of vesicle fusion ^20^. **4**) Optical assessment of vesicle pool dynamics was performed using Synapto-pHluorin-based imaging, inclusive of both pharmacological and genetic mechanistic dissection ^19^. **5**) Presynaptic calcium measurements were performed ^18^. **6)** Presynaptic capacitance measurements were performed ^18^. **7)** Multi-photon microscopy was used to assess live spine dynamics ^20^. **8**) 3D volumetric electron microscopy identified ultrastructural correlates of PHP termed ‘structural PHP’, inclusive of mechanistic pharmacological and genetic analyses ^19,20^. **9)** A comprehensive mechanistic dissection of PHP has been achieved including genome-scale forward genetics and genetic demonstration of mechanistic conservation across species and synapses ^16,19,21^. At the intersection of these diverse measures and experiments, a picture of PHP emerges at individual synapses that would otherwise be invisible to the limited tools of slice electrophysiology alone.

Here, we present new experimental evidence in support of hippocampal PHP. We replicate, and extend our previously published work, demonstrating how and why our assays are optimized. We emphasize the importance of the composition of the internal patch electrode solution. We identify a role for postsynaptic sub-threshold activity during PHP induction. We replicate and optimize an experimental protocol introduced by Nicoll and colleagues (Dou et al. 2026 BioRxiv) ^22^ and demonstrate that, once optimized, this protocol reveals robust PHP expression in adult mouse hippocampus. We directly compare our slice methodologies with those of Nicoll and colleagues using transmission electron microscopy, highlighting the importance of slice methods that have been optimized for use in adult brain. Finally, we demonstrate a negative impact of persistent, widespread axonal stimulation on the expression of PHP in adult brain slice. In conclusion, we reaffirm robust PHP in adult hippocampus (area CA1, *stratum oriens* ^19,20^) and, by extension, cerebellum ^18^. In so doing, we provide clear methodological and technical guidance for future studies of homeostasis in adult brain tissue.

## RESULTS

### Assay optimizations: Robust PHP expression during continuous patch-clamp recordings

As indicated above, the experimental evidence in support of PHP is extensive ^17–20^. Partial AMPA receptor antagonism was achieved using GYKI-53655, perampanel or JNJ55511118 across numerous experiments ^20^. One of these experiments involved washing-on GYKI and observing the rate of induction of PHP by continuously recording from a pyramidal neuron in adult hippocampal area CA1 (*stratum oriens*). Here, we replicate this experiment and extend our original findings (Figure 1A, D).

**Figure 1.**
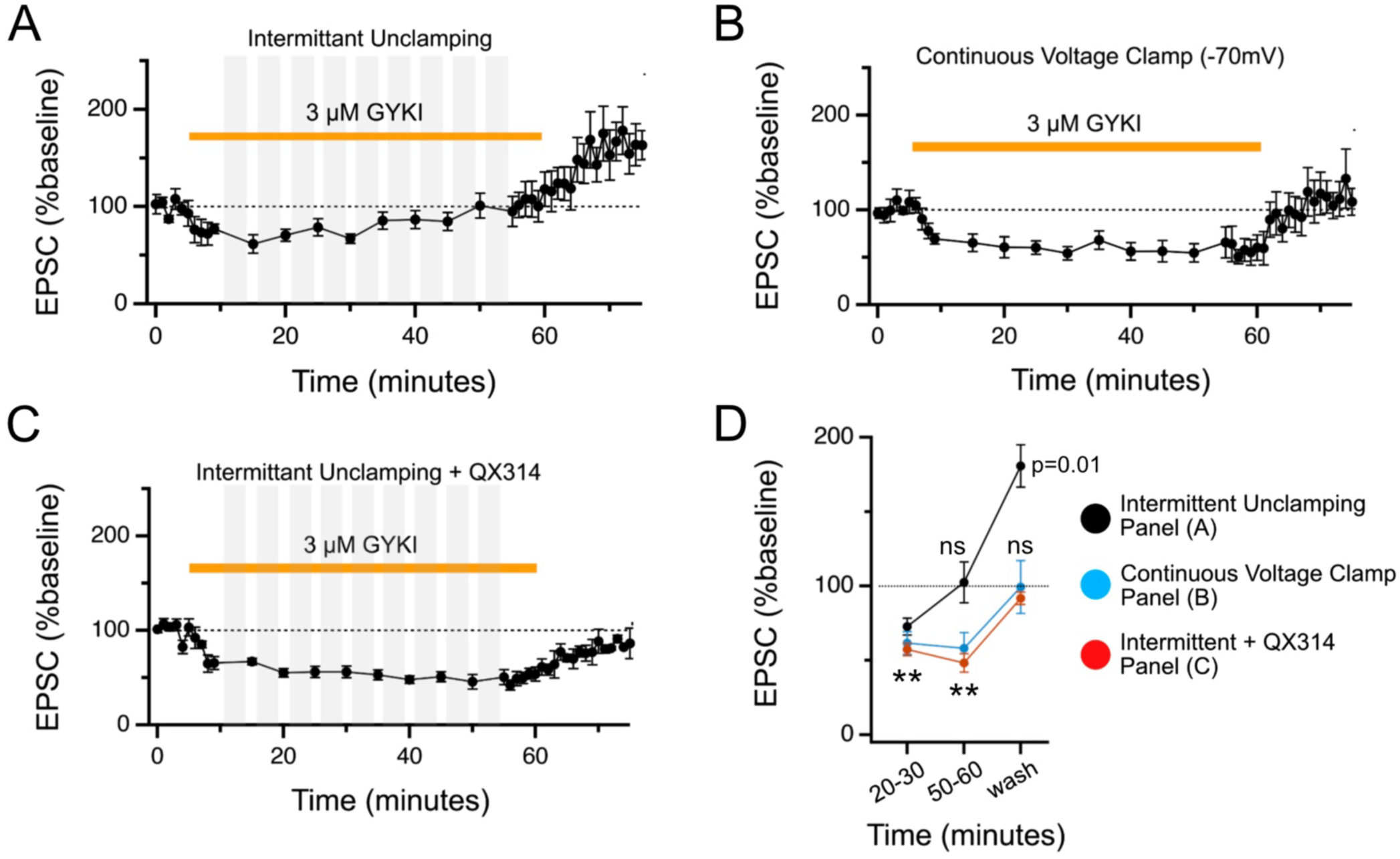
Robust PHP expression in adult hippocampus. **A)** Chronic recording from a pyramidal neuron (CA1) with stimulation (theta glass electrode, see methods) in *stratum oriens*. Gray shading indicates periods when the cell was unclamped. Orange bar indicates presence of GYKI (concentration indicated). GYKI washout begins at the end of orange bar. EPSC amplitudes normalized to baseline. N=4 animals, n=7 cells. **B)** Experiment as in (A) with continuous voltage clamp, i.e. without intermittent unclamping. N=6 animals, n=7 cells. **C)** Data as in (A) except QX314 is included in the patch electrode internal solution. N=5 animals, n=7 cells. **D)** Quantification of data in a single graph for experiments in A-C. Colored spheres indicate experimental conditions. Repeated measures One-Way ANOVA.

We first elaborate upon our experimental design and methodology, highlighting aspects of our work that we consider particularly important for experimental replication. First, we use a potassium-based internal patch electrode solution that lacks toxins such as cesium, spermine and QX314. These toxins block ion channels intracellularly and facilitate voltage clamp control of membrane potential (see METHODS) ^23–25^. Given the limited mechanistic understanding of hippocampal PHP, it is reasonable to assume that postsynaptic ion channels could participate (see next section), as they do in other forms of plasticity ^26–28^. Second, neurons are selected for patch clamp recording as far below the surface of the slice as possible while patching under visual control. We have previously shown that the integrity of experimental preparations can influence PHP in other systems ^29^, and we reason that by targeting neurons deeper within the slice we maximize the integrity of sampled synapses and minimize the negative impact of the sectioned neuronal processes along the cut surface. The health and integrity of cells, dendrites and synapses residing in the middle of our slices was confirmed by serial section transmission electron microscopy at time points that parallel our electrophysiological assessment of PHP (Chipman et al., 2022 ^20^ ; see also below). Third, small theta-glass electrodes are used with a barrel diameter of ∼1 µm, limiting tissue disruption during electrode placement. Finally, sub-blocking concentrations of GYKI are used, typically 3-5 µM, calibrated to achieve an approximate 50% reduction in stimulus-evoked unitary release event amplitude ^19,20^.

Next, we emphasize the importance of drug wash-off at the end of long patch clamp electrophysiology experiments. GYKI is a reversible, non-competitive allosteric AMPAR antagonist that can be readily washed from cells and out of brain slice^18,20,30–32^. When PHP is induced, GYKI wash-off results in EPSC amplitudes that over-shoot baseline amplitudes recorded prior to application of GYKI (Figure 1A) ^18,20^. This is diagnostic for PHP. By contrast, when PHP fails, then GYKI washout reveals EPSC amplitudes that simply return to baseline (Figure 1B,C). Indeed, PHP failure can only be distinguished from degraded cell health if EPSCs are found to return to baseline following GYKI washout. It is notable, therefore, that when Nicoll and colleagues perform GYKI washout, EPSC amplitudes either fail to show any recovery, or EPSC amplitudes continue to decline (see Dou et al., Figure 1) ^22^. Typically, when washout of a reversible antagonist fails, it reflects permanent, deleterious effects of an experiment on cell and tissue health.

### Assay optimization: Constant voltage clamp blocks PHP expression

Here, we provide new experimental data in support of one of our key methodological innovations. During our original PHP experiments, we included periods when the neuron was unclamped, thereby allowing the resting membrane potential to fluctuate. This was an attempt to minimally impact normal cellular physiology, allowing the system to express homeostasis. We reasoned that voltage fluctuation is also an essential component during other forms of activity-dependent plasticity ^33^.

We now provide experimental evidence to support our assertion that postsynaptic, sub-threshold activity is an important parameter for robust PHP induction. We repeated the experiments in Figure 1A under conditions of constant voltage clamp (using a potassium-based internal patch solution lacking ion channel toxins). We demonstrate that this causes PHP to fail (Figure 1B, D). Once again, please note that the absence of PHP is associated with a return of EPSC amplitudes precisely to baseline following GYKI wash-off. This is a clear indication that the cell, synapses and slice tissue remain healthy at the conclusion of our experiments. We are uncertain precisely why constant voltage clamp causes PHP failure. However, we note that low-voltage membrane changes can have profound effects on neurophysiology ^34,35^.

### Assay optimization: QX314 in the patch electrode blocks PHP expression

As stated above, our experiments to continuously monitor PHP induction (published and present) employ a potassium-based internal patch electrode solution that lacks cesium, spermine and QX314, all of which antagonize ion channels. Here, we directly demonstrate the importance of this assay optimization. We demonstrate that PHP can be robustly induced using a potassium gluconate-based internal solution lacking QX314 (Figure 1A). Next, we demonstrate that PHP fails when QX314 is included in the patch electrode (Figure 1C, D). It remains unknown whether QX314 antagonizes a postsynaptic channel necessary for PHP induction, or whether the activity of QX314 indirectly interferes with other postsynaptic processes necessary for PHP. By extension, it remains unknown whether the other toxins typically utilized in patch electrodes (cesium and spermine) might also adversely affect PHP. Finally, it should be emphasized that once PHP has been induced by application of GYKI, it can be subsequently assessed with an electrode containing cesium and QX314. We note that Nicoll and colleagues include QX314, spermine and cesium in all their patch-clamp experiments ^22^.

### Assay optimizations: Robust PHP using sequential two cell patch recordings

Nicoll and colleagues perform an experiment in which two cells are sequentially patched-clamped and synaptic transmission is assessed at a common set of inputs. In brief, the first cell establishes baseline stimulation conditions and is used to document the effects of GYKI wash-on. A second cell is patched at the end of the wash-on period (60 min of GYKI application) and is used to assess whether PHP has been induced.

We acknowledge that this new approach by Nicoll and colleagues can be used to circumvent the above issues regarding patch-electrode internal solution composition during persistent voltage clamp experiments. Therefore, we repeated their experiments using a sequential, two-cell experimental design. However, our experiments are further optimized in important ways. First, we use slice protocols optimized for adult brain slice (see below for further validation). Second, we eliminate stimulation during GYKI application as an unnecessary cellular disruption (see below for further validation). Third, we instituted an additional period of extended GYKI wash-out to confirm cell health. During this wash-out, PHP decays to baseline and EPSC amplitudes return to baseline. This is clear evidence that synapse and cell health have been maintained, consistent with experimental results from cerebellum ^18^.

In our experiment (Figure 2A), a recording from Cell1, prior to GYKI application, establishes baseline EPSC amplitudes (Figure 2B, C). The Cell1 patch electrode is then carefully removed. After 40 minutes of subsequent GYKI wash-on and incubation (note: no stimulation is performed that might alter slice physiology) a second cell (adjacent to the first) is patched (Cell2). Initial spontaneous and evoked amplitudes in Cell2 reveal the effects of GYKI (Figure 2B, C). First, we note that evoked amplitudes are less affected following 40min of GYKI incubation compared to spontaneous event amplitudes, consistent with the induction of PHP. We then wash GYKI out of the slice while maintaining the Cell2 patch recording. We demonstrate that washout restores spontaneous amplitudes of Cell2 toward the baseline amplitudes observed in Cell1 (Figure 2B) while the EPSC amplitudes of Cell2 significantly overshoot the baseline established in Cell1 (Figure 2C). This is diagnostic of PHP expression at Cell2. Finally, quantal content can be estimated by comparing Cell2 (post GYKI washout) with Cell1 because spontaneous EPSC amplitudes are nearly equivalent. This analysis demonstrates a significant ∼50% increase in estimated quantal content in Cell2. This is diagnostic of PHP expression (Figure 2D).

**Figure 2.**
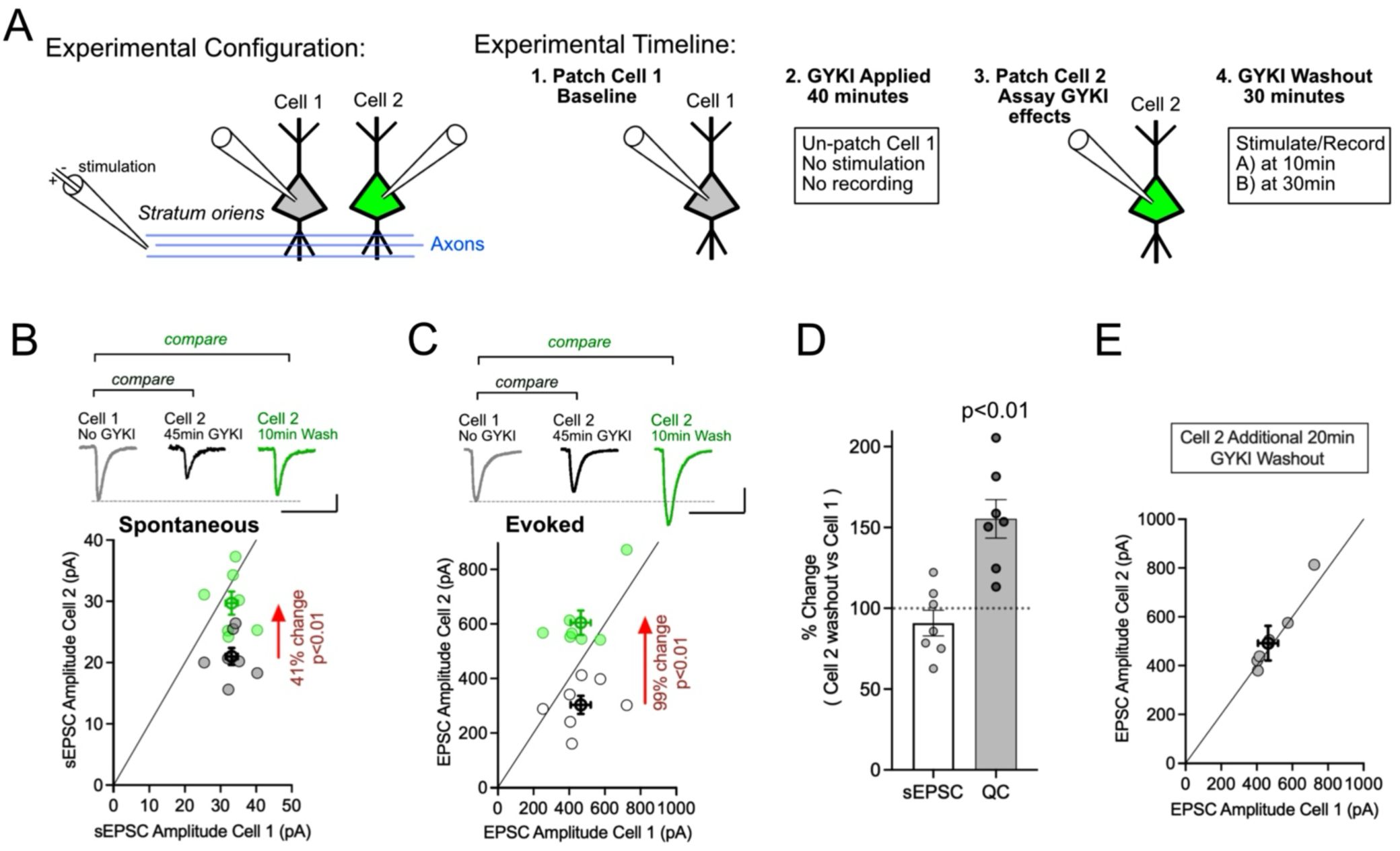
Robust PHP expression in adult hippocampus using sequential patching paradigm. **A)** Experimental design. Colors only indicate Cell1 and Cell2 for the purpose of clarity. **B)** Spontaneous miniature EPSC amplitudes for experiment outlined in (A). Cell2 washout results in a significant restoration of sEPSC amplitudes toward baseline values in Cell1. Average values are shown as symbols with thicker lines (+/- SEM). Paired t-test, two-way. N= 4 animals. All individual recordings shown. Scale 10pA, 50ms. **C)** Evoked EPSC amplitudes for identical cells shown in A and B. Cell2 washout results in a significant overshoot of EPSC amplitudes compared to values in Cell1, above the line of unity, diagnostic of PHP. Average values are shown as symbols with thicker lines (+/- SEM). Paired t-test, two-way. N= 4 animals. All individual recordings shown. Scale 200pA, 50ms. **D)** Estimated sEPSC amplitude and quantal content comparing Cell2 to Cell1 following 10min GYKI washout, normalized to values recorded in Cell1. **E)** Evoked EPSC amplitudes in Cell2 following 30min total GYKI washout, compared to Cell1. Values reside on the line of unity. Average values (+/- SEM).

Our version of the two-cell assay also includes an additional, <u>essential</u> control regarding cell and tissue health. We continued recording from Cell2 for an additional 20 minutes following GYKI washout (30 min total). EPSC amplitudes return to values identical to those observed in Cell1. This mirrors previously published data showing that PHP decays (∼30-90min) when GYKI is removed ^18^. The fact that EPSCs in Cell2 match those in Cell1 after 90 minutes of recording demonstrates that our slices remained healthy throughout our experiments (Figure 2E) (affirmed at the ultrastructural level, see below).

### Assay optimization: Improved adult brain slice health likely facilitates robust PHP expression

In our experiments, both published and current, we employ optimized protocols developed over the past decade for the adequate preservation of adult (>P35) mammalian (including human) brain tissue ^36–46^. By contrast, Nicoll and colleagues use a slicing protocol commonly employed to obtain tissue from younger animals (P20-35) that is based on use of high-sucrose containing ACSF for slicing and recovery (see Methods). Here, we directly compared these two protocols using transmission electron microscopy, assessing multiple time points that parallel critical moments during electrophysiological PHP assessment.

First, several published optimizations for adult slice preparation are highlighted including the use of vascular perfusion of ice-cold cutting solutions immediately prior to brain removal, optimization of slice orientation, the adjustment of cutting solution osmolarity (305 mOsm +/- 5 mOsm) and close monitoring of pH (7.4 +/- 0.5). Vascular perfusion should enable oxygenated and pH balanced ACSF solutions to access the synaptic neuropil, mitigating potentially damaging impact of anoxic blood. The optimization of slice orientation preserves the integrity of CA1 pyramidal neuron dendritic arbors from the *stratum oriens* to the *stratum lacunosum moleculare*. The proper adjustment of osmolarity and pH minimizes cell swelling and death. All our efforts in this regard are an attempt to enable the highest possible tissue quality. To confirm our tissue quality, we routinely verify tissue integrity in slices that are fixed and analyzed using transmission electron microscopy. Once again, because we are studying a homeostatic process, we reason that every effort needs to be made to optimize cell health and integrity.

In new experiments presented here, we prepared acute slices from adult brain according to our previously published, optimized protocol (as per above). Our new EM data precisely (quantitatively, see Table 1 - and qualitatively) parallel our published data, revealing well-preserved cell bodies, dendrites, axons and synapses at all time points (Figure 3A). Subsynaptic features, including docked vesicles, postsynaptic densities and other active zone features are clear. The mitochondria appear healthy with intact cristae, microtubules are apparent, and both the endoplasmic reticulum and spine apparatus can be readily detected. It should be noted that all EM images are acquired at the center of the acute slice, demonstrating robust preservation of synapse morphology at the site of cell patching and synapse function. We would also like to emphasize that all our published 3D EM volumes are from brain slices that are prepared identically to slices used for our electrophysiology experiments, allowing direct comparison between functional assessments and tissue integrity (see Chipman et al., 2022 ^20^; 2025 ^19^)

**Figure 3.**
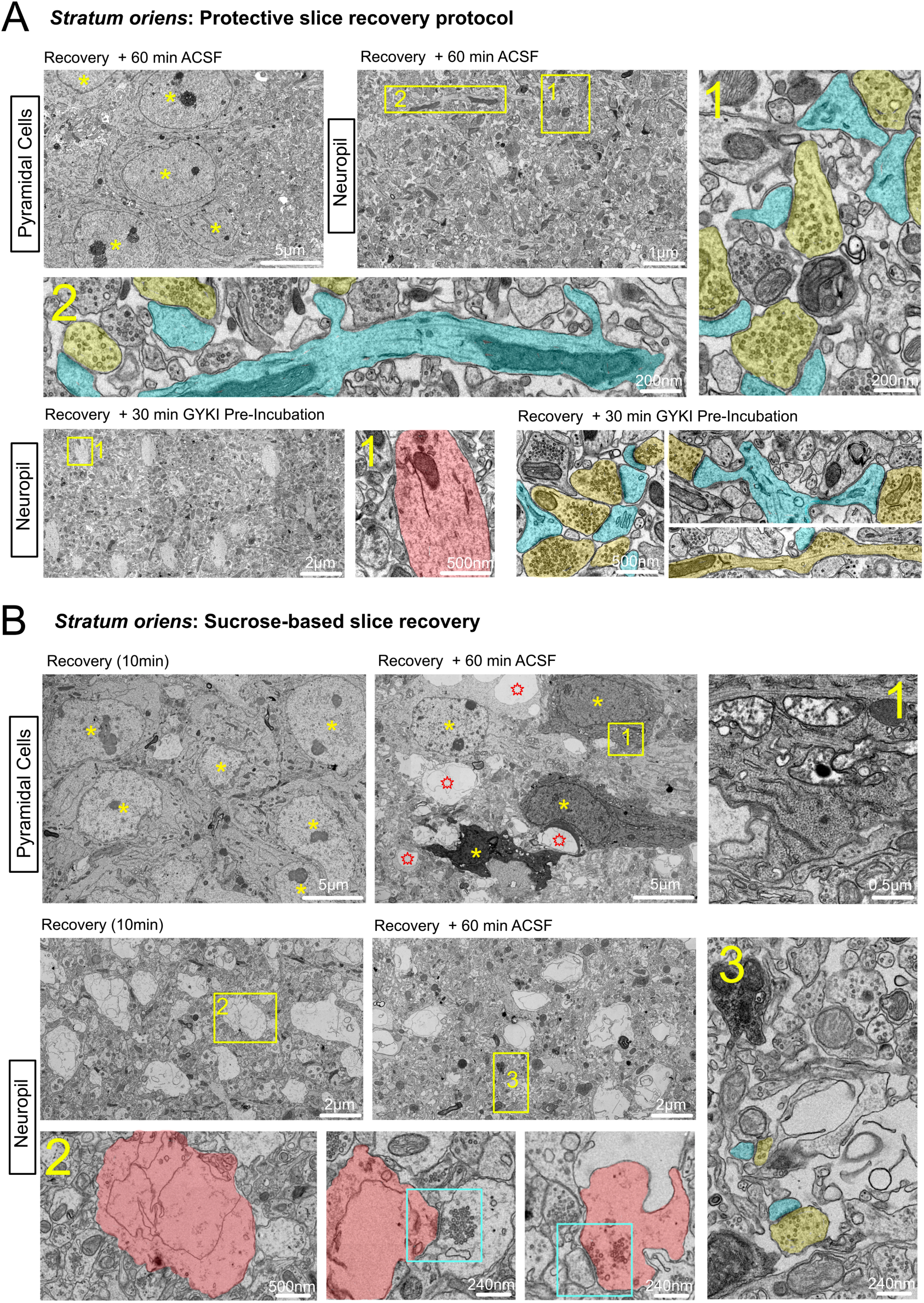
Cell and synapse degeneration associated with sucrose-based brain slice protocol. **A)** Data from multiple brain slices prepared using protective slice recovery protocols deployed by the Davis laboratory in this and in prior publications ^19,20^. The image labeled ‘Pyramidal Cells’ is a montage taken in the cell body layer of hippocampus (CA1) at a late time point (Recovery plus 60 min incubation in ASCF) paralleling the precise time point when PHP is typically assessed in brain slice electrophysiologically. At this time point, cell bodies (yellow asterisks) and neuropil appear healthy. The image labeled ‘Neuropil’ is a montage from *stratum oriens* at the indicated time point. Yellow box #1 and the associated inset highlights synapse density and integrity (yellow is presynaptic and blue is postsynaptic). Yellow box #2 and the associated inset show a healthy dendrite (blue) with healthy mitochondria surrounded by synapses (yellow presynaptic, blue postsynaptic). The bottom row of images show data from a slice prepared for electrophysiology including GYKI incubation. All images are from neuropil in *stratum oriens*. The presence of GYKI does not alter tissue, cell or synapse integrity. Yellow box #1 highlights the profile of a large dendrite with a robust cytoskeleton, mitochondria and ER. Images at right highlight synapse integrity and density. See Table 1 for quantification of synapse density in these and additional EM montages. **B)** Data from multiple brain slices prepared using the sucrose-based brain slice methodology employed by Dou et al., (2026) ^22^. The row of images (left to right) indicated as “Pyramidal Cells” are montages taken in the cell body layer of hippocampus (CA1). Two time points are shown as indicated. Following 10min of recovery, cell bodies appear healthy (yellow asterisks). Following an additional 60min in ACSF, cell bodies show evidence of necrosis (black) and degeneration (darkened cytoplasm). The inset (yellow box #1 and higher magnification at right) highlights severely compromised mitochondria within the neuronal cell body of a neuron with darkened cytoplasm. In the second row of images, two low-mag images (labeled 10min and +60min respectively) indicated as “Neuropil” are montages of hippocampus (CA1) *stratum oriens* at indicated time points post slicing. Large voids are present throughout the montage, indicated by the region in yellow box #2, shown at higher magnification below (inset 2). The red color indicates the void as a distended, membrane bound structure. At right, two additional high-magnification images reveal remnants of a synapse in which the red colored void is postsynaptic (middle, see blue box) and a synapse in which the red void is presynaptic (right, see blue box). Inset number 3 highlights an area of neuropil and the number of synapses contained within that area, presynaptic is yellow and postsynaptic is blue. Degenerating cellular material is evident in the dark intracellular objects and empty membranes.

**TABLE 1.** Adverse consequences of sucrose-based slice preparation in adult brain.

| Condition: tissue preparation for 2D transmission EM | Active zone count / 25μm<sup>2</sup> | Active zone density | % Change to Control | Source of Data |
| --- | --- | --- | --- | --- |
| Published Control <sup>1</sup> | 240 | 9.6 / μm<sup>2</sup> | Control | Chipman et al., 2022 |
| Adult optimized: Recovery <sup>2</sup> | 243 | 9.7 / μm<sup>2</sup> | Plus 1% | This manuscript |
| Adult optimized: Recovery + 60min <sup>3</sup> | 225 | 9.0 / μm<sup>2</sup> | Minus 6% | This manuscript |
| Sucrose-based: Recovery <sup>4</sup> | 96 | 3.8 / μm<sup>2</sup> | Control | This manuscript |
| Sucrose-based: Recovery + 60min <sup>5</sup> | 77 | 3.1 / μm<sup>2</sup> | Minus 20% | This manuscript |
| Sucrose-based: Recovery + 60min + stim <sup>6</sup> | 53 | 2.1 / μm<sup>2</sup> | Minus 45% | This manuscript |

**Legend:** Transmission electron microscopy (TEM) analysis of adult hippocampal CA1. Animal age was P90 for Published Control and all other samples were P55. Montages (25 x 25 μm, single plane) were analyzed for each sample, visually identifying every active zone. 'Percent Change to Control' (column 4) compares quantification within groups defined by sample preparation.

Next, we prepared slices from adult brain using the sucrose-based slice protocol (Figure 3B). We find that pyramidal neuron cell bodies initially appear healthy (10min slice recovery). However, after a further 60min of slice incubation in an electrophysiology recording chamber perfused with oxygenated ACSF the tissue shows clear and remarkable degradation (Figure 3B). Cell bodies are dark and necrotic. Mitochondria are severely compromised, lacking intact cristae. In addition, we demonstrate that widespread, adverse effects occur immediately in the neuropil of the *stratum oriens* and become worse with time. We document widespread dendrite and axonal destruction (revealed as voids). These voids are distended membranes without evidence of intracellular cytoskeleton, mitochondria or internal membranes (e.g. endoplasmic reticulum). When comparing the effects of different slice protocols and considering the impact on the expression of PHP, the quantification of synapse density (Table 1) is particularly revealing. The sucrose slicing protocol results in diminish synapse density by more than half at the very outset (10 min slice recovery), and there is a further decrease in synapse density of 45% over a time course (60min) that parallels the timecourse of electrophysiological experiments performed by Dou et al., (2026) ^22^.

We acknowledge that our electron microscopy experiments do not directly assess the integrity of tissue prepared by the Nicoll laboratory, or any other laboratory. Yet, our experiment was guided by the details of published methodology ^22^ and our data agree with prior studies that emphasize the benefits of adult-optimized slicing protocols ^36–46^. Thus, we suggest that poor tissue integrity and progressive degeneration are potential factors contributing to failed PHP in the experiments performed by Nicoll and colleagues. For example, progressive synapse degeneration could reasonably contribute to failed GYKI washout (Dou et al., Figure 1) ^22^. Furthermore, PHP is associated with physical expansion of the active zone and postsynaptic spine (Chipman et al., 2022; 2025). These changes could reasonably be constrained or prevented under conditions of synapse loss. Finally, degeneration-associated cell stress could reasonably interfere with signaling necessary for the induction and expression of PHP ^9^.

### Assay optimization: Continuous low-frequency stimulation disrupts PHP expression

In our published work ^19,20^, as well as experiments presented here (Figures 1 and 2), we avoid continuous electrical stimulation during the period of GYKI incubation. Nerve stimulation, particularly with a bi-polar electrode such as deployed by Nicoll and colleagues ^22^, is not a neutral event. Large numbers of axons and dendrites are typically recruited with each stimulation, influencing the physiology of axons, synapses and dendrites in the region of stimulation. Here, we demonstrate that continuous electrical stimulation at a rate of 0.033Hz (1stim / 30 seconds) is sufficient to block PHP induction with deleterious effects that persist following GYKI washout.

We repeated the sequential two-cell assay from Figure 2 with one change. We add a second stimulation electrode, a standard concentric bipolar electrode, positioned slightly further away from our patched cells on the same side as our small theta-glass electrode (Figure 4A). In brief, initial EPSC amplitudes in Cell1 are established with theta-glass electrode stimulation. GYKI is washed into the slice and bipolar electrode stimulation is initiated at a rate of 0.033Hz for the 40min duration of GYKI incubation. Cell2 is patched and the effects of GYKI are determined using theta-glass electrode stimulation. GYKI is washed out and EPSCs are again determined for Cell2 using theta glass stimulation at two time points (10min and 30min) (Figure 4A). Bipolar electrode stimulus intensity was set at the half-maximal synaptic recruitment intensity (∼0.2mA), achieving an average EPSC amplitude (average = 890pA) in Cell1 that is ∼230% larger than that achieved by the theta-glass electrode stimulus (average = 390pA).

**Figure 4.**
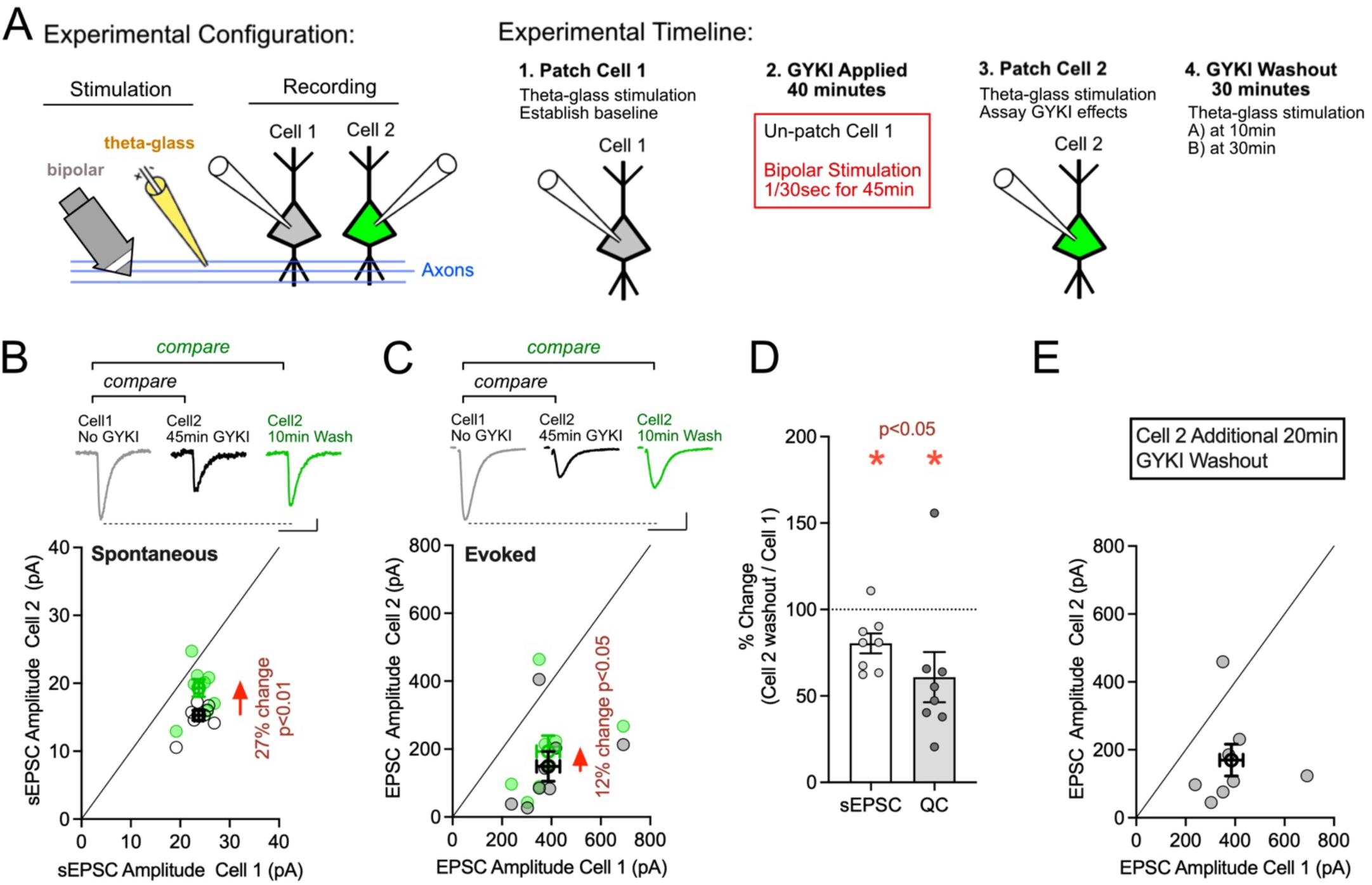
Continuous stimulation blocks the induction of PHP. **A)** Experimental design. Colors only indicate Cell1 and Cell2 for the purpose of clarity. **(B)** Spontaneous miniature EPSC amplitudes for experiment outlined in (A). GYKI suppresses sEPSC amplitudes to 63% of Cell 1 (p<0.001, paired t-test). Partial recovery after 10-minute washout is observed (27%; paired t-test). Scale bar = 10 pA, 50 ms. **(C)** Evoked EPSC amplitudes for identical cells shown in A and B. Evoked EPSCs remain suppressed after washout with only 12% recovery (p=0.01, paired t-test). Average values are shown as symbols with thicker lines (+/- SEM). Scale bar = 200 pA, 50 ms. **D**) Estimated sEPSC amplitude and quantal content comparing Cell2 to Cell1 following 10min GYKI washout, normalized to values recorded in Cell1. **(E)** Evoked EPSC amplitudes in Cell2 following 30min total GYKI washout, compared to Cell1. Evoked responses remain suppressed below unity line (p=0.0156, Wilcoxon test). Data points are individual cells values, and bold points are mean ± SEM. N = 8 paired recordings (16 cells) from 4 mice.

The critical comparisons for assessment of PHP are at the 10min and 30min GYKI washout timepoints. When we incorporate constant stimulation during GYKI application, we still observe a small, significant recovery of spontaneous event amplitudes in Cell2 (27%), moving toward baseline recorded in Cell1 (Figure 4B). However, EPSC amplitudes show only a 12% recovery and do not approach baseline values in Cell1 (Figure 4C). Estimation of quantal content demonstrates a complete absence of PHP (Figure 4D). Indeed, quantal content falls significantly below baseline in Cell1, consistent with a persistent deleterious effect of chronic stimulation. This conclusion is supported by the failure of EPSCs to recover after an additional 20min of GYKI washout (compare Figure 4E with Figure 2E). Thus, continuous low frequency stimulation appears deleterious and is sufficient to block PHP expression. Our experimental conditions are designed to parallel those used by Nicoll and colleagues, referring both to their 2-cell sequential patch clamp experiment and their experiment using an extracellular electrode to record field potentials (notably reported for recordings in *stratum radiatum* though they write that identical results were found in *stratum oriens* but those data are not shown). Thus, the use of chronic low-frequency stimulation is another variable that could reasonably contribute to the failure of Nicoll and colleagues to document PHP.

## DISCUSSION

We reaffirm that PHP is robustly expressed in adult hippocampus. Our findings are consistent with, and extend, our published work that documented a required function for NMDARs and an evolutionarily conserved role for Semaphorin/Integrin signaling as an underlying molecular mechanism ^19,20^. In so doing, we highlight the essential importance of assay optimizations that were employed in our previously published work and provide additional experimental guidance for future studies of homeostasis in adult brain tissue.

Homeostasis is a means by which cells respond to a perturbation and sustain normal functionality. This is the original definition of what is now considered a pillar of modern cellular and organismal physiology ^9,47^. The many forms of homeostasis (PHP included) are intimately and necessarily tied to cell health. If cell health is sufficiently compromised, then normal cell function degrades, cellular stress dominates and homeostasis fails. This is widely considered a pathophysiological progression underlying neurological and psychiatric disease ^48–50^. Thus, a prerequisite for studying PHP is to ensure adequate cell/tissue health.

### The basis for assay optimization

Experiments can fail for many reasons including the unintended incorporation of methodological or procedural errors. Our approach has been to incorporate diverse techniques (electrophysiological, optical and electron microscopy) to create a foundation for studying PHP in the adult mammalian brain. At the same time, we recognize known limitations associated with each approach and this has driven our efforts at assay optimization. While patch-clamp electrophysiology in acute slice is a powerful means to reveal the features of neuron and synapse function, the technique is limited in the ability to precisely control the voltage at the active zone in the spine head ^23,51^. As such, direct optical assessment of vesicle release provides independent confirmation of PHP expression ^19,20^. Likewise, ultrastructural changes in vesicle pools are well matched to both electrophysiology and optical data.

#### Discovery of experimental variables relevant to PHP

Homeostasis, by definition, is a process that is intimately connected to the healthy physiology of a cell or tissue. As such, the incorporation of toxins into an experiment design needs to be considered as a confound that could disrupt homeostasis. Here, we formally demonstrate that continuous voltage clamp and/or inclusion of QX314 in the postsynaptic cell will block PHP induction. We have yet to test whether inclusion of spermine or cesium, as used by Nicoll and colleagues ^22^, also disrupt PHP expression. Our published methods are also clear regarding the temperature at which experiments are performed (34°C). It remains unknown whether key processes necessary for the rapid induction of PHP are temperature dependent, but it is not unreasonable to consider this as an important factor.

Finally, we demonstrate that continuous stimulation during GYKI application, using a bipolar electrode commonly deployed in hippocampal electrophysiology ^22^, is sufficient to block the induction of PHP. We note that PHP is an activity-independent phenomenon in every system where it has been studied to date. It is not unreasonable, therefore, to discover that imposing activity via the stimulation of axons, synapses and dendrites is deleterious to PHP induction. Indeed, this observation could represent an important point of intersection between the mechanisms of activity-dependent plasticity and activity-independent PHP. Accordingly, the induction of learning related plasticity would not be opposed by simultaneous recruitment of PHP.

### Mechanistic dissection underscores the existence of PHP

Despite our focus on experimental paradigm, it is important not to lose sight of our previously published mechanistic advances. The required function of NMDA receptors during PHP was a serendipitous discovery during early electrophysiological experiments, subsequently validated using EM as well as optical analyses of synaptic transmission (these were performed in an independent laboratory). The required function of NMDA receptors was further validated with a dual patch clamp assessment in single slices. Two adjacent cells were patched, one electrode included MK801 and the other did not. In the presence of perampanel, the MK801 infused cell reported a smaller average EPSC, consistent with PHP failure ^20^. Again, these data support the conclusion that NMDA receptors are necessary for PHP induction and/or expression.

In our recent publication, the secreted signal Sema3a was shown to be required for PHP across three experimental modalities (electrophysiology, optical analyses and EM), inclusive of both loss of function and rescue experiments ^19^. The demonstration that these mechanisms did not substantively alter baseline transmission but blocked PHP provides powerful confirmation of the phenomenon. Finally, we would like to emphasize that mechanistic discovery confers additional confidence in the phenomenon of PHP for a straightforward reason. When testing whether a genetic mutation, interfering protein or drug blocks PHP, our experiments are performed blind. As a result, each mechanistic iteration becomes a validation that PHP is robustly expressed by healthy tissue, regardless of whether or not the gene or drug is found to alter expression of PHP.

### Regarding previously published studies of AMPAR antagonism and NMDA receptor currents

Nicoll and colleagues reference several publications in which AMPARs are manipulated genetically, either by Gria subunit deletion or knockout of proteins that localize to the postsynaptic density, including associated signalling molecules ^52–54^. In the cited work, changes in NMDAR currents were not observed. First, we note that changes in NMDAR currents can be separated from expression of PHP as shown in Chipman et al., 2022. Notably, triple *<u>heterozygous</u>* depletion of the Gria1, Gria2 and Gria3 AMPA receptor subunits does not lead to enhanced NMDAR-mediated neurotransmission, though other features of PHP are observed, including alterations in calcium-dependence of vesicle fusion and short-term plasticity ^20^. Furthermore, it should be noted that enhanced NMDA currents are observed following loss of Gria4 in cerebellum ^55^. Thus, while NMDARs are necessary for PHP induction, enhanced NMDA currents are a *correlate* of PHP expression. Second, many of the accessory, signaling, and scaffolding molecules highlighted by the Nicoll group could reasonably impact the expression of PHP when deleted ^56–58^. Finally, the determination of PHP in a genetic mutant requires the application of experiments outlined in our prior publications which determine how a given mutation differentially affects unitary and evoked EPSC amplitudes (see next section). Since PHP was not the focus of that prior work, it is understandable that it was not documented.

### The importance of quantifying both unitary and evoked synaptic events for PHP assessment

In our previously published results ^19,20^, the assessment of PHP is informed by the relative changes that occur to quantal and evoked synaptic events. Evidence of PHP emerges when evoked EPSC amplitudes return to baseline despite continued effects of GYKI on quantal amplitudes (please see Figure 2, above). Furthermore, quantal amplitudes also inform regarding the stability of receptor antagonism over time, an important indication of cell and tissue health. It is, therefore, notable that spontaneous event amplitudes are either not measured or not reported in the data presented by Nicoll and colleagues ^22^.

### Complexity and unknowns

Homeostasis is a complex biological process, whether considering synaptic transmission, neuronal excitability or cellular proteostasis ^9,10,12,47^. 25 years of electrophysiology-based forward genetic screens have begun to reveal molecular mechanisms of PHP, but these data only scratch the surface ^29,59,60^. We do not know how partial AMPAR antagonism initiates PHP. We do not understand the required function(s) of NMDA receptors, or how the rapid induction of PHP is converted into long-term expression. We do not yet understand the link with inhibitory synaptic plasticity or the full impact of PHP on neural circuit function. These are among many unknowns that are reasons to experiment and explore.

## RESOURCE AVAILABILITY

### Lead contact

Further information and requests for resources and reagents should be directed to and will be fulfilled by the Lead Contact, Graeme W. Davis.

### Materials Availability

Upon final publication, all unique/stable reagents generated in this study are available from the lead contact without restriction.

### Data and code availability

- This paper does not report original code.
- All data reported in this paper will be shared by the lead contact upon request.

## ACKNOWLEDGEMENTS

Supported by NINDS grant R35NS137247 to GWD.

## AUTHOR CONTRIBUTIONS

PHC, RDF, FJR, UL: design, data collection, analyses, interpretation, co-writing and editing manuscript.

GWD: funding acquisition, project design, data analysis interpretation, writing and editing manuscript.

## METHODS

### EXPERIMENTAL MODEL AND SUBJECT DETAILS

#### Mouse lines and knockout genetics

Male and female *C57BL6/J* (IMSR_JAX:000664) were obtained as adults (6-10 weeks old) from The Jackson Laboratory. All procedures were performed in accordance with UCSF (protocol # AN108729-02B) IACUC guidelines.

### Acute brain slice preparation

#### Slice protocol: Davis laboratory

Male and female *C57BL6/J* between the ages of ∼7-14 weeks were used for the preparation of acute slices as described in previously published protocols ^19,20,36,61^. Briefly, mice were deeply anesthetized with isoflurane and transcardially perfused with cutting aCSF solution containing (in mM): 93 N-methyl D-glucamine, 2.5 KCl, 1.2 NaH_2_PO_4_, 30 NaHCO_3_, 20 HEPES, 20 glucose, 5 Na ascorbate, 2 thiourea, 3 sodium pyruvate, 12 N-acetyl L-cysteine, 10 MgSO_4_, 0.5 CaCl_2_, pH adjusted to 7.4 with HCl and bubbled with 95% O_2_ / 5% CO_2_, ∼300 mOsm. Blood was first cleared in near room temperature cutting solution to prevent vascular constriction, then switched to ice-cold cutting solution to slow cellular metabolism and reduce dissection-associated inflammation. Brains were extracted and fixed to the cutting stage with cyanoacrylate-based tissue adhesive positioned at a ∼30-40° angle from horizontal along the rostral/caudal axis using a 4% agar block. 350 μm transverse hippocampal sections of were obtained in ice-cold cutting aCSF with a ceramic blade (Cadence blades #EFINZ10), and a Leica VT1200 vibrating microtome. Hemispheres were separated and small cuts were made near the CA2/CA1 border to prevent recurrent activity during axonal stimulation in electrophysiology experiments. Slices were incubated for 12 minutes in cutting aCSF warmed to 34°C, then placed in holding aCSF solution containing (in mM) 81.2 NaCl, 2.5 KCl, 1.2 NaH_2_PO_4_, 30 NaHCO_3_, 20 HEPES, 20 D-glucose, 5 Na ascorbate, 2 thiourea, 3 sodium pyruvate, 12 N-acetyl L-cysteine, 2 MgSO_4_, 2 CaCl_2_, pH 7.4, bubbled with 95% O_2_ / 5% CO_2_, ∼300 mOsm at room temperature (∼20°C) for at least 1 hour and up to 8 hours until used in experiments.

#### Slice protocol reported in (Dou et al., 2026) ^22^

The methodology presented in the study by Nicoll and colleagues does not directly report the composition of their slicing solution. Neither does the work referenced in their results section. We interpret the Nicoll group’s reference to “standard, long established methods” as a sucrose-based slicing approach, typically used for the preparation of acute brain slices from neonatal and juvenile rodents, which was detailed in a recent Nicoll laboratory publication ^62^. This solution contains (in mM) 2.5 KCl, 7 MgSO_4_, 1.25NaH_2_PO_4_, 25 NaHCO_3_, 7 glucose, 210 sucrose, 1.3 ascorbic acid. The remainder of the protocol was followed as described in the manuscript by Nicoll and colleagues ^22^.

### Electrophysiology

Davis lab acute slice protocols as described above were used for electrophysiology experiments. Whole-cell patch clamp recordings were obtained from CA1 pyramidal neurons using an Olympus BX51W1 microscope equipped with IR-DIC optics and a motorized stage. Pyramidal neurons were visually identified by their large, almond-shaped cell bodies and position within the pyramidal cell layer. Experiments were carried out using Multiclamp 700B amplifiers and Clampex10.7 acquisition software (Molecular Devices). Analysis was performed using Clampfit10.7 and MiniAnalysis software. Patch pipettes (borosilicate glass, OD 1.5mm, ID 0.86mm, tip resistance 2-4 MΩ) were pulled using a Sutter P-97 micropipette puller. Slices were constantly perfused with recording aCSF at 1.5-2mL/min containing (in mM) 119 NaCl, 2.5 KCl, 1.3 NaH_2_PO_4_, 26 NaHCO_3_, 1 MgCl_2_, 2 CaCl_2_, 20 D-glucose and 0.5 Na ascorbate pH 7.4, bubbled with 95% O_2_ / 5% CO_2_, ∼295-305 mOsm and maintained at 32-34°C using an in-line heater (Harvard Instruments). Picrotoxin (100 µM; Tocris #1128) was added to the recording aCSF to isolate glutamatergic synaptic transmission. We used two internal pipette solutions. One contained (in mM) 130 CsMeSO_3_, 8 NaCl, 4 Mg-ATP, 0.3 Na-GTP, 0.5 EGTA, 10 HEPES, pH 7.3, 5 QX314-bromide (Tocris #2555), ∼290-295 mOsm. The other contain 142 K-gluconate, 10 HEPES, 1 EGTA, 2.5 Mg_2_Cl, 4 Mg_2_-ATP, 0.3Na_3_-GTP, 10 Na-phosphocreatine, pH 7.3, 290-295 mOsm. Patch solutions were allowed >10 minutes to equilibrate through the cell before experiments were performed. Pipette series resistances were ∼20 MΩ and were compensated by ∼30-60% in some experiments to achieve R_s_ values of <10 MΩ (see below). Experiments in which uncompensated R_s_ was >30 MΩ or changed by >20% were discarded.

#### Continuous EPSC monitoring experiments

Experiments in which EPSCs were continually monitored throughout the duration the experiment were performed in near physiological concentrations of calcium and magnesium (2mM [Ca^2+^]_e_/1mM [Mg^2+^]_e_) and in the presence of picrotoxin (100 µM). Whole cell patch configuration was achieved using K-gluconate-based internal solutions as described above. Stimulation electrodes were constructed from theta glass (borosilicate glass, OD 1.5 mm, ID 1.00 mm, SEP 0.2 mm, tip diameter ∼ 1-3 µm) pulled to a tip diameter of ∼1µm and positioned within ∼200 µm of the patched neuron in the middle of the *stratum oriens*. Tungsten wire electrodes were placed in either barrel of the theta glass pipettes and attached to the positive and negative poles of a stimulus isolation unit (A.M.PI. ISO-Flex; Jerusalem, Israel). Stable EPSCs were achieved by carefully adjusting the position of stimulation electrodes during a pre-experiment baseline sampling period. Once stable responses were achieved, the position and strength of the stimulation electrode was fixed and not further altered for the duration of the experiment.

#### Sequential Patch experiments

This experiment used cesium-based internal solution (see above) and theta-glass stimulation electrode pulled to a tip diameter of ∼1 µm and placed as per above. Cell 1 was patched and a stable baseline of evoked EPSCs and spontaneous sEPSCs were established. Once stable responses were achieved, the position and strength of the stimulation electrode was fixed and unaltered for the duration of the experiment. Following baseline, the recording pipette was removed while the stimulation pipette remained fixed. GYKI was bath-applied for 40 minutes. Then, Cell 2 was patched, allowed to equilibrate for 10min followed by acquisition of both spontaneous and evoked EPSCs. GYKI was subsequently washed out for 10 minutes and both spontaneous and evoked EPSCs were sampled again. After an additional 20 minutes washout, both spontaneous and evoked EPSCs were sampled for a final time.

#### Continuous bipolar stimulation

A concentric bipolar microelectrode (2–3 μm inner tip diameter, 126 μm outer conductor width, ∼200 kΩ impedance; WPI cat# PTM3CC02INS) was positioned in the *stratum oriens* adjacent to the theta-glass electrode, slightly more distal compared to the recorded cell. Concentric electrode stimulus amplitude was titrated to evoke a large EPSC, approximately half-maximal synaptic recruitment and without substantial waveform broadening. Concentric stimulation intensity averaged 0.2 mA (± 0.1 mA SD).

### Electron Microscopy

Acute brain slices were prepared as described above for electrophysiology experiments, either according to the methodology of Nicoll and colleagues ^22^ or the Davis laboratory ^19,20^. After recovery and any additional post-recovery experimental conditions outlined above, slices were treated identically. Slices were fixed by immersion up to 1 hour in 2% glutaraldehyde in 0.1M Na-cacodylate buffer, pH 7.4 at room temperature (RT) followed by overnight at 4° C. Fixed slices were then post-fixed with 1% OsO_4_/1.5% KFe(CN)_6_/0.1 M Na-cacodylate for 1 hr at RT, followed by 1% OsO_4_/0.1M Na-cacodylate for 1 hr at RT, *en bloc* staining in 5% uranyl acetate in water for 1 hr at RT, dehydration, infiltration and polymerization in Eponate 12 resin (Ted Pella, Inc., Redding, CA). Sections (40nm thickness) of the *stratum oriens* were cut with a Leica UC7 ultramicrotome using a Diatome diamond knife, picked up on Pioloform coated slot grids and stained with uranyl acetate and Sato’s lead. Sections were imaged with an FEI Tecnai T12 TEM at 120 kV using a Gatan Rio 4k x 4k camera. Montages (25 x 25 µm, single plane) were imaged using SerialEM and aligned with TrakEM2/Fiji. Modeling and analysis were performed with IMOD.

### QUANTIFICATION AND STATISTICAL ANALYSIS

All statistical analysis were performed using Origin Pro 9, Prism 8, or Igor Pro 8. When means are shown, error bars indicate standard error. Parametric or non-parametric statistical analyses were performed when data were normally distributed and when normality could be rejected, respectively. Statistical tests used are indicated in figure legends inclusive of sample size and animal number as appropriate.

## REFERENCES

1. Davis, G.W., and Goodman, C.S. (1998). Synapse-specific control of synaptic efficacy at the terminals of a single neuron. Nature 392, 82–86. 10.1038/32176.

2. MacLean, J.N., Zhang, Y., Johnson, B.R., and Harris-Warrick, R.M. (2003). Activity-independent homeostasis in rhythmically active neurons. Neuron 37, 109–120.

3. Davis, G.W. (2006). HOMEOSTATIC CONTROL OF NEURAL ACTIVITY: From Phenomenology to Molecular Design. Annu Rev Neurosci 29, 307–323. 10.1146/annurev.neuro.28.061604.135751.

4. Turrigiano, G., Abbott, L.F., and Marder, E. (1994). Activity-dependent changes in the intrinsic properties of cultured neurons. Science (New York, NY) 264, 974–977. 10.1126/science.8178157.

5. Turrigiano, G.G., Leslie, K.R., Desai, N.S., Rutherford, L.C., and Nelson, S.B. (1998). Activity-dependent scaling of quantal amplitude in neocortical neurons. Nature 391, 892–896. 10.1038/36103.

6. Béïque, J.C., Na, Y., Kuhl, D., Worley, P.F., and Huganir, R.L. (2011). Arc-dependent synapse-specific homeostatic plasticity. Proceedings of the National Academy of Sciences 108, 816–821. 10.1073/pnas.1017914108/-/dcsupplemental.

7. Keck, T., Keller, G.B., Jacobsen, R.I., Eysel, U.T., Bonhoeffer, T., and Hübener, M. (2013). Synaptic Scaling and Homeostatic Plasticity in the Mouse Visual Cortex In Vivo. Neuron 80, 327–334. 10.1016/j.neuron.2013.08.018.

8. Driscoll, L.N., Pettit, N.L., Minderer, M., Chettih, S.N., and Harvey, C.D. (2017). Dynamic Reorganization of Neuronal Activity Patterns in Parietal Cortex. Cell 170, 986–999.e16. 10.1016/j.cell.2017.07.021.

9. Davis, G.W. (2013). Homeostatic Signaling and the Stabilization of Neural Function. Neuron 80, 718–728. 10.1016/j.neuron.2013.09.044.

10. Vitureira, N., Letellier, M., and Goda, Y. (2012). Homeostatic synaptic plasticity: from single synapses to neural circuits. Current opinion in neurobiology 22, 516–521. 10.1016/j.conb.2011.09.006.

11. Deeg, K.E., and Aizenman, C.D. (2011). Sensory modality–specific homeostatic plasticity in the developing optic tectum. Nat. Neurosci. 14, 548–550. 10.1038/nn.2772.

12. Wen, W., and Turrigiano, G.G. (2024). Keeping Your Brain in Balance: Homeostatic Regulation of Network Function. Annu. Rev. Neurosci. 47, 41–61. 10.1146/annurev-neuro-092523-110001.

13. Orr, B.O., Hauswirth, A.G., Celona, B., Fetter, R.D., Zunino, G., Kvon, E.Z., Zhu, Y., Pennacchio, L.A., Black, B.L., and Davis, G.W. (2020). Presynaptic Homeostasis Opposes Disease Progression in Mouse Models of ALS-Like Degeneration: Evidence for Homeostatic Neuroprotection. Neuron 107, 95–111.e6. 10.1016/j.neuron.2020.04.009.

14. Wang, X., McIntosh, J.M., and Rich, M.M. (2018). Muscle Nicotinic Acetylcholine Receptors May Mediate Trans-Synaptic Signaling at the Mouse Neuromuscular Junction. The Journal of neuroscience 38, 1725–1736. 10.1523/jneurosci.1789-17.2018.

15. Wang, X., Pinter, M.J., and Rich, M.M. (2016). Reversible Recruitment of a Homeostatic Reserve Pool of Synaptic Vesicles Underlies Rapid Homeostatic Plasticity of Quantal Content. Journal of Neuroscience 36, 828–836. 10.1523/jneurosci.3786-15.2016.

16. Orr, B.O., Fetter, R.D., and Davis, G.W. (2022). Activation and expansion of presynaptic signaling foci drives presynaptic homeostatic plasticity. Neuron. 10.1016/j.neuron.2022.08.016.

17. Cull-Candy, S.G., Miledi, R., Trautmann, A., and Uchitel, O.D. (1980). On the release of transmitter at normal, myasthenia gravis and myasthenic syndrome affected human end-plates. J Physiology 299, 621–638. 10.1113/jphysiol.1980.sp013145.

18. Delvendahl, I., Kita, K., and Müller, M. (2019). Rapid and sustained homeostatic control of presynaptic exocytosis at a central synapse. Proceedings of the National Academy of Sciences 21, 201909675. 10.1073/pnas.1909675116.

19. Chipman, P.H., Lee, U., Orr, B.O., Fetter, R.D., and Davis, G.W. (2025). A unifying mechanism for presynaptic homeostatic plasticity at mammalian peripheral and central synapses. Neuron. 10.1016/j.neuron.2025.05.030.

20. Chipman, P.H., Fetter, R.D., Panzera, L.C., Bergerson, S.J., Karmelic, D., Yokoyama, S., Hoppa, M.B., and Davis, G.W. (2022). NMDAR-dependent presynaptic homeostasis in adult hippocampus: Synapse growth and cross-modal inhibitory plasticity. Neuron. 10.1016/j.neuron.2022.08.014.

21. Orr, B.O., Fetter, R.D., and Davis, G.W. (2017). Retrograde semaphorin-plexin signalling drives homeostatic synaptic plasticity. Nature Publishing Group 550, 109–113. 10.1038/nature24017.

22. Dou, T., Zhang, J., Hong, Y., Chen, X., and Nicoll, R.A. (2026). NMDA receptor-dependent presynaptic homeostatic plasticity? BioRxiv [preprint], 10.64898/2026.02.28.708706.

23. Williams, S.R., and Mitchell, S.J. (2008). Direct measurement of somatic voltage clamp errors in central neurons. Nature neuroscience 11, 790–798. 10.1038/nn.2137.

24. Talbot, M.J., and Sayer, R.J. (1996). Intracellular QX-314 inhibits calcium currents in hippocampal CA1 pyramidal neurons. Journal of neurophysiology 76, 2120–2124. 10.1152/jn.1996.76.3.2120.

25. Bowie, D., and Mayer, M.L. (1995). Inward rectification of both AMPA and kainate subtype glutamate receptors generated by polyamine-mediated ion channel block. Neuron 15, 453–462. 10.1016/0896-6273(95)90049-7.

26. Chen, H.-X., Otmakhov, N., and Lisman, J. (1999). Requirements for LTP Induction by Pairing in Hippocampal CA1 Pyramidal Cells. J. Neurophysiol. 82, 526–532. 10.1152/jn.1999.82.2.526.

27. Watanabe, S., Hoffman, D.A., Migliore, M., and Johnston, D. (2002). Dendritic K+ channels contribute to spike-timing dependent long-term potentiation in hippocampal pyramidal neurons. Proc. Natl. Acad. Sci. 99, 8366–8371. 10.1073/pnas.122210599.

28. Kullmann, D.M., Perkei, D.J., Manabe, T., and Nicoll, R.A. (1992). Ca2+ Entry via postsynaptic voltage-sensitive Ca2+ channels can transiently potentiate excitatory synaptic transmission in the hippocampus. Neuron 9, 1175–1183. 10.1016/0896-6273(92)90075-o.

29. Frank, C.A., Kennedy, M.J., Goold, C.P., Marek, K.W., and Davis, G.W. (2006). Mechanisms Underlying the Rapid Induction and Sustained Expression of Synaptic Homeostasis. Neuron 52, 663–677. 10.1016/j.neuron.2006.09.029.

30. Paternain, A.V., Morales, M., and Lerma, J. (1995). Selective antagonism of AMPA receptors unmasks kainate receptor-mediated responses in hippocampal neurons. Neuron 14, 185–189. 10.1016/0896-6273(95)90253-8.

31. Balannik, V., Menniti, F.S., Paternain, A.V., Lerma, J., and Stern-Bach, Y. (2005). Molecular Mechanism of AMPA Receptor Noncompetitive Antagonism. Neuron 48, 279–288. 10.1016/j.neuron.2005.09.024.

32. Yelshanskaya, M.V., Singh, A.K., Sampson, J.M., Narangoda, C., Kurnikova, M., and Sobolevsky, A.I. (2016). Structural Bases of Noncompetitive Inhibition of AMPA-Subtype Ionotropic Glutamate Receptors by Antiepileptic Drugs. Neuron 91, 1305–1315. 10.1016/j.neuron.2016.08.012.

33. Magee, J.C., and Johnston, D. (1997). A Synaptically Controlled, Associative Signal for Hebbian Plasticity in Hippocampal Neurons. Science 275, 209–213. 10.1126/science.275.5297.209.

34. Awatramani, G.B., Price, G.D., and Trussell, L.O. (2005). Modulation of Transmitter Release by Presynaptic Resting Potential and Background Calcium Levels. Neuron 48, 109–121. 10.1016/j.neuron.2005.08.038.

35. Christie, J.M., Chiu, D.N., and Jahr, C.E. (2010). Ca2+-dependent enhancement of release by subthreshold somatic depolarization. Nature Publishing Group 14, 62–68. 10.1038/nn.2718.

36. Ting, J.T., Daigle, T.L., Chen, Q., and Feng, G. (2014). Acute Brain Slice Methods for Adult and Aging Animals: Application of Targeted Patch Clamp Analysis and Optogenetics. In Patch-Clamp Methods and Protocols., pp. 221–242. 10.1007/978-1-4939-1096-0_14.

37. Ting, J.T., Lee, B.R., Chong, P., Soler-Llavina, G., Cobbs, C., Koch, C., Zeng, H., and Lein, E. (2018). Preparation of Acute Brain Slices Using an Optimized <em>N</em>-Methyl-D-glucamine Protective Recovery Method. Journal of Visualized Experiments, 1–13. 10.3791/53825.

38. Cadwell, C.R., Palasantza, A., Jiang, X., Berens, P., Deng, Q., Yilmaz, M., Reimer, J., Shen, S., Bethge, M., Tolias, K.F., et al. (2015). Electrophysiological, transcriptomic and morphologic profiling of single neurons using Patch-seq. Nature biotechnology 34, 199–203. 10.1038/nbt.3445.

39. Jiang, X., Shen, S., Cadwell, C.R., Berens, P., Sinz, F., Ecker, A.S., Patel, S., and Tolias, A.S. (2015). Principles of connectivity among morphologically defined cell types in adult neocortex. Science (New York, N.Y.) 350, aac9462–aac9462. 10.1126/science.aac9462.

40. Hendricks, W.D., Chen, Y., Bensen, A.L., Westbrook, G.L., and Schnell, E. (2017). Short-Term Depression of Sprouted Mossy Fiber Synapses from Adult-Born Granule Cells. J. Neurosci. 37, 5722–5735. 10.1523/jneurosci.0761-17.2017.

41. Kalmbach, B.E., Buchin, A., Long, B., Close, J., Nandi, A., Miller, J.A., Bakken, T.E., Hodge, R.D., Chong, P., Frates, R. de, et al. (2018). h-Channels Contribute to Divergent Intrinsic Membrane Properties of Supragranular Pyramidal Neurons in Human versus Mouse Cerebral Cortex. Neuron 100, 1194–1208.e5. 10.1016/j.neuron.2018.10.012.

42. Tasic, B., Yao, Z., Graybuck, L.T., Smith, K.A., Nguyen, T.N., Bertagnolli, D., Goldy, J., Garren, E., Economo, M.N., Viswanathan, S., et al. (2018). Shared and distinct transcriptomic cell types across neocortical areas. Nature 563, 72–78. 10.1038/s41586-018-0654-5.

43. Asrican, B., Wooten, J., Li, Y.-D., Quintanilla, L., Zhang, F., Wander, C., Bao, H., Yeh, C.-Y., Luo, Y.-J., Olsen, R., et al. (2020). Neuropeptides Modulate Local Astrocytes to Regulate Adult Hippocampal Neural Stem Cells. Neuron 108, 349–366.e6. 10.1016/j.neuron.2020.07.039.

44. Bakken, T.E., Jorstad, N.L., Hu, Q., Lake, B.B., Tian, W., Kalmbach, B.E., Crow, M., Hodge, R.D., Krienen, F.M., Sorensen, S.A., et al. (2021). Comparative cellular analysis of motor cortex in human, marmoset and mouse. Nature 598, 111–119. 10.1038/s41586-021-03465-8.

45. Conner, J.M., Bohannon, A., Igarashi, M., Taniguchi, J., Baltar, N., and Azim, E. (2021). Modulation of tactile feedback for the execution of dexterous movement. Science 374, 316–323. 10.1126/science.abh1123.

46. Soden, M.E., Yee, J.X., and Zweifel, L.S. (2023). Circuit coordination of opposing neuropeptide and neurotransmitter signals. Nature 619, 332–337. 10.1038/s41586-023-06246-7.

47. Billman, G.E. (2020). Homeostasis: The Underappreciated and Far Too Often Ignored Central Organizing Principle of Physiology. Front. Physiol. 11, 200. 10.3389/fphys.2020.00200.

48. Bourgeron, T. (2015). From the genetic architecture to synaptic plasticity in autism spectrum disorder. Nat Rev Neurosci 16, 551–563. 10.1038/nrn3992.

49. Frere, S., and Slutsky, I. (2018). Alzheimer’s Disease: From Firing Instability to Homeostasis Network Collapse. Neuron 97, 32–58. 10.1016/j.neuron.2017.11.028.

50. Radulescu, C.I., Doostdar, N., Zabouri, N., Melgosa-Ecenarro, L., Wang, X., Sadeh, S., Pavlidi, P., Airey, J., Kopanitsa, M., Clopath, C., et al. (2023). Age-related dysregulation of homeostatic control in neuronal microcircuits. Nat. Neurosci. 26, 2158–2170. 10.1038/s41593-023-01451-z.

51. Beaulieu-Laroche, L., and Harnett, M.T. (2018). Dendritic Spines Prevent Synaptic Voltage Clamp. Neuron 97, 75–82.e3. 10.1016/j.neuron.2017.11.016.

52. Díaz-Alonso, J., Sun, Y.J., Granger, A.J., Levy, J.M., Blankenship, S.M., and Nicoll, R.A. (2017). Subunit-specific role for the amino-terminal domain of AMPA receptors in synaptic targeting. Proc. Natl. Acad. Sci. 114, 7136–7141. 10.1073/pnas.1707472114.

53. Lu, W., Shi, Y., Jackson, A.C., Bjorgan, K., During, M.J., Sprengel, R., Seeburg, P.H., and Nicoll, R.A. (2009). Subunit Composition of Synaptic AMPA Receptors Revealed by a Single-Cell Genetic Approach. Neuron 62, 254–268. 10.1016/j.neuron.2009.02.027.

54. Watson, J.F., Ho, H., and Greger, I.H. (2017). Synaptic transmission and plasticity require AMPA receptor anchoring via its N-terminal domain. eLife 6, e23024. 10.7554/elife.23024.

55. Kita, K., Albergaria, C., Machado, A.S., Carey, M.R., Müller, M., and Delvendahl, I. (2021). GluA4 facilitates cerebellar expansion coding and enables associative memory formation. eLife 10, e65152. 10.7554/elife.65152.

56. Haghighi, A.P., McCabe, B.D., Fetter, R.D., Palmer, J.E., Hom, S., and Goodman, C.S. (2003). Retrograde Control of Synaptic Transmission by Postsynaptic CaMKII at the Drosophila Neuromuscular Junction. Neuron 39, 255–267. 10.1016/s0896-6273(03)00427-6.

57. Goel, P., Li, X., and Dickman, D. (2017). Disparate Postsynaptic Induction Mechanisms Ultimately Converge to Drive the Retrograde Enhancement of Presynaptic Efficacy. CellReports 21, 2339–2347. 10.1016/j.celrep.2017.10.116.

58. Qiu, C., Perry, S., Chen, C., Chen, J., Zhuang, J., Han, Y., Goel, P., and Dickman, D. (2025). Nonionic signaling rapidly remodels postsynaptic DLG to induce retrograde homeostatic plasticity. Proc. Natl. Acad. Sci. 122, e2502997122. 10.1073/pnas.2502997122.

59. Dickman, D.K., and Davis, G.W. (2009). The schizophrenia susceptibility gene dysbindin controls synaptic homeostasis. Science (New York, N.Y.) 326, 1127–1130. 10.1126/science.1179685.

60. Davis, G.W., and Goodman, C.S. (1998). Genetic analysis of synaptic development and plasticity: homeostatic regulation of synaptic efficacy. Curr Opin Neurobiol 8, 149–156. 10.1016/s0959-4388(98)80018-4.

61. Chipman, P.H., Fung, C.C.A., Fernandez, A.P., Sawant, A., Tedoldi, A., Kawai, A., Gautam, S.G., Kurosawa, M., Abe, M., Sakimura, K., et al. (2021). Astrocyte GluN2C NMDA receptors control basal synaptic strengths of hippocampal CA1 pyramidal neurons in the stratum radiatum. Elife 10, e70818. 10.7554/elife.70818.

62. Díaz-Alonso, J., Morishita, W., Incontro, S., Simms, J., Holtzman, J., Gill, M., Mucke, L., Malenka, R.C., and Nicoll, R.A. (2020). Long-term potentiation is independent of the C-tail of the GluA1 AMPA receptor subunit. eLife 9, e58042. 10.7554/elife.58042.

